# A microRNA profile in *Fmr1* knockout mice reveals microRNA expression alterations with possible roles in fragile X syndrome

**DOI:** 10.1101/002071

**Authors:** Ting Liu, Rui-Ping Wan, Ling-Jia Tang, Shu-Jing Liu, Hai-Jun Li, Qi-Hua Zhao, Wei-Ping Liao, Xiao-Fang Sun, Yong-Hong Yi, Yue-Sheng Long

**Affiliations:** Key Laboratory of Neurogenetics and Channelopathies of Guangdong Province and the Ministry of Education of China, Institute of Neuroscience and the Second Affiliated Hospital of Guangzhou Medical University, Guangzhou 510260, China; Key Laboratory for Major Obstetric Diseases of Guangdong Province and Key laboratory of Reproduction and Genetics of Guangdong Higher Education Institutes, Guangzhou Medical University, Guangzhou 510150, China

**Keywords:** microRNA, transcriptome profiling, FMRP, fragile X syndrome

## Abstract

Fragile X syndrome (FXS), a common form of inherited mental retardation, is caused by a loss of expression of the fragile X mental retardation protein (FMRP). FMRP is involved in brain functions by interacting with mRNAs and microRNAs (miRNAs) that selectively control gene expression at translational level. However, little is known about the role of FMRP in regulating miRNA expression. Here, we found a development-dependant dynamic expression of *Fmr1* mRNA (encoding FMRP) in mouse hippocampus with a small peak at postnatal day 7 (P7). MiRNA microarray analysis showed that the levels of 38 miRNAs showed a significant increase with about 15∼250 folds and the levels of 26 miRNAs showed a significant decrease with only about 2∼4 folds in the hippocampus of P7 *Fmr1* KO mice. Q-PCR assay showed that 9 of the most increased miRNAs (>100 folds in microarrays) were increased about 40∼70 folds and their pre-miRNAs were increased about 5∼10 folds, but no significant difference in their pri-miRNA levels was observed, suggesting a role of FMRP in regulating miRNA processing from pri-miRNA to pre-miRNA. We further demonstrated that a set of protein-coding mRNAs, potentially targeted by the 9 miRNAs were down-regulated in the hippocampus of *Fmr1* KO mice. Finally, luciferase assays demonstrated that miR-34b, miR-340, miR-148a could down-regulate the reporter gene expression by interacting with the *Met* 3′ UTR. Taken together these findings suggest that the miRNA expression alterations resulted from the absence of FMRP might contribute to molecular pathology of FXS.

## Introduction

The *FMR1* gene encodes a fragile X mental retardation protein (FMRP) that plays a critical role in brain development and is involved in synaptic plasticity (De Boulle et al. 1993; Huber et al. 2002). Abnormal *FMR1* gene expression results in several neurodevelopmental and neurological disorders including fragile X syndrome (FXS), a common form of inherited intellectual disability (Bailey et al. 2001; Hessl et al. 2005; Jacquemont et al. 2007; O’Donnell and Warren 2002; Verkerk et al. 1991). Most FXS cases are caused by unstable expansion of the CGG trinucleotide repeat in the promoter region of the *FMR1* gene that leads to a low level of FMRP by translational suppression (Feng et al. 1995; Kremer et al. 1991). A few cases are caused by deleting part or all of the *FMR1* gene (Hirst et al. 1995; Trottier et al. 1994), or by point mutations that lead to a change in one of the amino acids in the gene (Chelly and Mandel 2001; Siomi et al. 1994; Zang et al. 2009).

FMRP, predominantly expressed in the brain, is involved in synaptic plasticity by transferring mRNAs from nucleus to cytoplasm, localizing dendritic mRNA, and regulating synaptic protein synthesis (Antar et al. 2005; Darnell and Klann 2013; Darnell et al. 2011). FMRP-regulated the productions of new proteins in response to synaptic activities at dendrites allow for the various physical changes and frequently-changing synaptic connections that are linked with the processes of learning and memory (Mercaldo et al. 2009; Sidorov et al. 2013). It has been demonstrated that FMRP can selectively bind mRNAs with some degree of sequence specificity (Ashley et al. 1993; Blackwell et al. 2010; Siomi, Choi, Siomi, Nussbaum and Dreyfuss 1994), and transfer mRNA from nucleus to cytoplasm (Eberhart et al. 1996; Feng et al. 1997). Generally, FMRP is predominantly localized in the cytoplasm and is occasionally present in the nucleus (Verheij et al. 1993). However, FMRP is also primarily present in the nucleus during a specific developmental period, implicating an important nuclear role in the brain development (Blonden et al. 2005).

FMRP is also involved in microRNA (miRNA) pathway by targeting with miRNAs or interacting with the Dicer and Argonaute proteins in the cytoplasm (Duan and Jin 2006; Jin et al. 2004), suggesting an important role of FMRP in miRNAs maturation. A recent study shows that FMRP-associated miRNAs regulate dendritic spine morphology and synaptic function which are important for the pathogenesis of FXS (Edbauer et al. 2010). One hypothesis is that a very low level of FMRP resulted from overexpansion of the CGG trinucleotide repeat in FXS patients could lead to abnormal expression of miRNAs that might be involved in the pathogenesis of FXS. It is unknown which specific miRNAs are regulated by FMRP during miRNA maturation. We speculate that the levels of some specific miRNAs might be altered by the lack of FMRP in *Fmr1* knockout (KO) mice, which could result in synaptic abnormalities. However, to our knowledge, there has no comprehensive analysis of the miRNA expression in *Fmr1* KO mice or in the brain tissues of FXS patients.

In the present study, we, for the first time, provided a differential genome-wide miRNA profile in the hippocampus of *Fmr1* KO mice comparing to that in the wildtype mice. In *Fmr1* KO mice, the number and the change extent of the up-regualted miRNAs were larger than those of the down-regulated miRNAs. We then demonstrated that the up-regulations of the miRNAs are resulted from miRNA processing of pri-miRNA into pre-miRNA, instead of miRNA gene transcription. This result indicates a new role of FMRP in the miRNA pathway. Furthermore, we demonstrated that expressions of the protein-coding mRNAs potentially targeted by those increased miRNAs were down-regulated in the *Fmr1* KO mice. Therefore, this study provides a differential miRNA profile in the *Fmr1* KO mice that is a model of FXS, and then suggests that the altered miRNAs could be used as potential biomarkers for the diagnosis of this disease.

## Results

### Dynamic transcription of *Fmr1* in the mouse hippocampus

Previous studies have shown that FMRP plays an important role in brain development (Lu et al. 2004; Michel et al. 2004). To determine the temporal expression pattern of *Fmr1* in the mouse hippocampus, we extracted total RNA from the mouse hippocampus at different stages from embryonic day 18 (E18) to postnatal day 60 (P60). We found that *Fmr1* mRNA transcripts could be detected at all stages using RT-PCR (**Figure 1A**). The qRT-PCR experiment showed that the relative mRNA expression levels (*Fmr1*/β-actin) at P1, P7 and P15 were about 0.1, 0.6 and 0.2 fold of that at E18, respectively. No significance was found between E18 and P60 (*P* > 0.05). Significant alterations (*P* < 0.001) in the relative mRNA expression levels were observed between two adjacent stages (**Figure 1B**). These data suggest a development-dependant dynamic expression of *Fmr1* in the mouse hippocampus.

**Figure 1.**
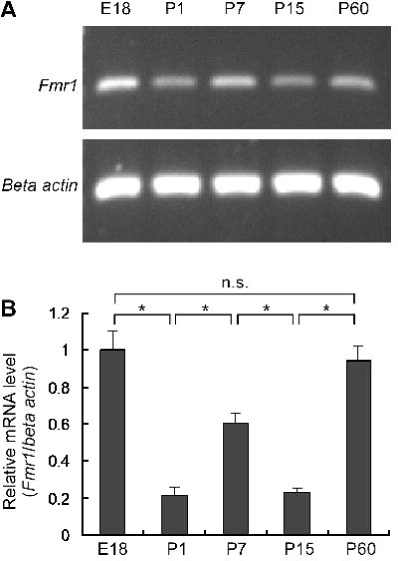
Dynamic expression of *Fmr1* mRNA in mouse hippocampus. (*A*) Detection of mouse *Fmr1* mRNA in the hippocampus by RT-PCR. E18 indicates embryonic day 18; P1, P7, P15 and P60 indicate postnatal day 1, 7, 15 and 60, respectively. (*B*) *Fmr1* expression at different developmental stages measured by qRT-PCR. The *Fmr1* mRNA levels were normalized to the mRNA levels of *β-actin*, a house-keep gene. The relative gene expression (*Fmr1/β-actin*) in E18 mice was normalized as “1”. \**P* < 0.001, n=9; n.s. represents no significance. All experiments were performed with a biological repeat of 3 individual mice and a technical replicate for three times.

### Altered miRNA expression in the hippocampus of *Fmr1* KO mice

we speculate that FMRP might be involved in miRNA expression since FMRP is a selective RNA-binding protein and is associated with miRNA pathway (Ashley, Wilkinson, Reines and Warren 1993; Duan and Jin 2006; Jin, Zarnescu, Ceman, Nakamoto, Mowrey, Jongens, Nelson, Moses and Warren 2004). However, a genome-wide miRNA profile in the *Fmr1* KO mice has not been reported. We initially determined that FMRP was not expressed in the hippocampus of *Fmr1* KO mice using Western blot analysis (**Figure 2A**). We determined the miRNA expression levels in the hippocampus of 3 *Fmr1* KO mice and 3 wildtype mice at P7 by hybridization of the array containing 1080 miRNAs. After normalization of control oligos, the differential expression of miRNAs between the KO mice and the wildtype mice was quantified using a phosphorimager (**Figure 2B**). Compared with the wildtype mice, 38 of the 1080 miRNAs in the KO mice showed up-regulated expression with *P* value < 0.01. Of them, 33 miRNAs had an increase with about 15∼250 folds (**Figure 2C**, **Table S1).** Twenty-six miRNAs in the KO mice showed down-regulated expression with a decrease of only about 2∼4 folds (**Figure 2C**, **Table S2**). According to change extent of the miRNA expression, we suggest that FMRP tends to negatively regulate miRNA expression. To further validate our results, 9 of the most up-regulated miRNAs with changes >100-fold were confirmed by qRT-PCR analysis (Table 1). We found that all these 9 miRNAs were increased about 40∼70 folds in the KO mice (**Figure 3A**).

### The up-regulated miRNAs in the hippocampus of *Fmr1* KO mice are resulted from miRNA processing of pri-miRNA into pre-miRNA

To demonstrate whether the increased miRNAs in the *Fmr1* KO mice are resulted from miRNA processing or miRNA transcription, we used qRT-PCR to quantify the pri-miRNA and pre-miRNA of the up-regulated miRNAs. **Figure 3B** showed that the relative levels (pre-miRNA/U6) of the 9 pre-miRNAs in the KO mice were about 5∼10 folds of that in the wildtype mice. However, no significant difference in the relative levels (pri-miRNA/U6) of the 9 pri-miRNAs was observed between the KO mice and the wildtype mice (**Figure 3C**). These data indicate that the up-regulated mature miRNAs in the *Fmr1* KO mice are resulted from miRNA processing of pri-miRNA into pre-miRNA, instead of miRNA transcription.

### Down-regulation of miRNA target genes in the hippocampus of *Fmr1* KO mice

We searched the potential target genes of the altered miRNAs using bioinformatic analysis. About 5000 genes were predicted as the targets for the 9 up-regulated miRNAs (**Table S3**). Of them, 392 genes were associated with neuronal function and 4484 genes were associated with other functions (**Figure 4A**, **Table S4**). Due to the role of FMRP in the brain, we paid more attention to the target genes that are associated with neuronal function. **Figure 4B** showed that most of the 392 genes had only one potential target for the 9 increased miRNAs and 6 genes had up to 6 targets. Using qRT-PCR analysis, we detected the mRNA transcripts of 9 target genes which are highly expressed in mouse hippocampus according to the public data from Allen Brain Atlas Database (**Figure S1**) and have 5 or 6 potential microRNA targets in their 3′ UTRs (**Table S5**). We found that the relative mRNA levels of the 9 genes in the hippocampus of the KO mice were all significantly lower than those in the wildtype mice (**Figure 5**), suggesting that the expression of the target genes were down-regulated in the *Fmr1* KO mice.

### Overexpression of miRNAs down-regulate the reporter gene expression through interacting with the *Met* 3′ UTR

We selected mouse *Met* gene as one example to identify the potential miRNA targets described above. Four miRNAs, miR-34b-5p, miR-340-5p, miR-101a and miR-148a-3p, were found to potentially target on the *Met* 3′ UTR with computer program prediction. Of them, miR-34b-5p had two potential targets (**Figure 6A**). To examine whether the 4 miRNAs regulate gene expression, we produced a serial of recombinant constructs psiCHECK2-mMet-F (F indicates different 3′-UTR fragments), which contain one cassette expressing synthetic *Renilla* luciferase (hRluc) and the other cassette expressing synthetic firefly luciferase (hluc+) that was used as an internal control reporter (**Figure 6B**). We produced a serial of GFP fusion pre-miRNA constructs pCMV-EGFP-pre-miRNA. Luciferase assay showed that the relative luciferase activities (hRluc/hluc+) of psiCHECK2-mMet-F1 cotransfected with pCMV-EGFP-miR-34b in N1E-115 cells and Neuro-2A cells were respectively only ∼70% and ∼50% of that cotransfected with pCMV-EGFP. The relative activities of psiCHECK2-mMet-F2 cotransfected with pCMV-EGFP-miR-340 in these two cell lines were respectively ∼75% and ∼66% of that cotransfected with pCMV-EGFP. The relative activities of psiCHECK2-mMet-F3 cotransfected with pCMV-EGFP-miR-148a in these two cell lines were respectively ∼73% and ∼63% of that cotransfected with pCMV-EGFP. However, no significant difference in luciferase activity was observed in the other transfections (**Figure 6C**). These data show that miR-34b, miR-340, miR-148a could down-regulate the reporter gene expression by interacting with the *Met* 3′ UTR.

## Discussion

In this study, we, for the first time, provide a genome-wide miRNA expression profile in the hippocampus of *Fmr1* KO mice using a miRNA microarray containing about 1000 miRNAs, revealing the altered expressions of a number of miRNAs in the KO mice comparing to that in the wildtype mice. By qRT-PCR analysis, we demonstrated that 9 of the up-regulated miRNA with over 100 folds were altered at both the mature miRNA level and the pre-miRNA level, but not at the pri-miRNA level, suggesting that the altered expression of these miRNAs should be resulted from miRNA processing of pri-miRNA into pre-miRNA, instead of miRNA gene transcription. Furthermore, we have also demonstrated that the expressions of the potentially targeted protein-coding genes by the 9 increased miRNAs were down-regulated in the *Fmr1* KO mice. The luciferase assay demonstrated that 3 of the 5 targets in the mouse *Met* 3′ UTR were functional in two cell lines. Together, our results suggest a FMRP-miRNAs-targets pathway that may play an important role in regulating brain development and synaptic plasticity.

**Figure 2.**
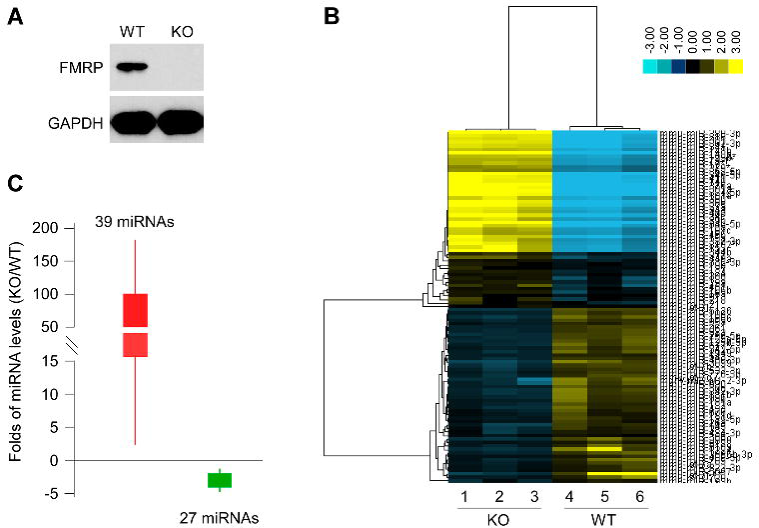
Differentially expressed miRNAs in the hippocampus of *Fmr1* knockout P7 mice. (*A*) Western blot analysis of FMRP expression in the hippocampus of Wild-type (WT) mouse and *Fmr1* knockout (KO) mouse. GAPDH expression was also detected for the loading control. (*B*) Hierarchical clustering of 77 differentially expressed miRNAs with *P* value < 0.05 (from Table S1 and Table S2) between 3 WT and 3 KO mice. Tissue samples are represented in the columns, and differentially expressed miRNAs are delineated in rows. Color bar shows that yellow indicates up-regulation and green represents down-regulation. KO samples and WT samples were clustered separately. (*C*) The changing folds of the 39 up-regulated miRNAs and the 27 down-regulated miRNAs with *P* value < 0.01 in the hippocampus of *Fmr1* KO mice.

**Figure 3.**
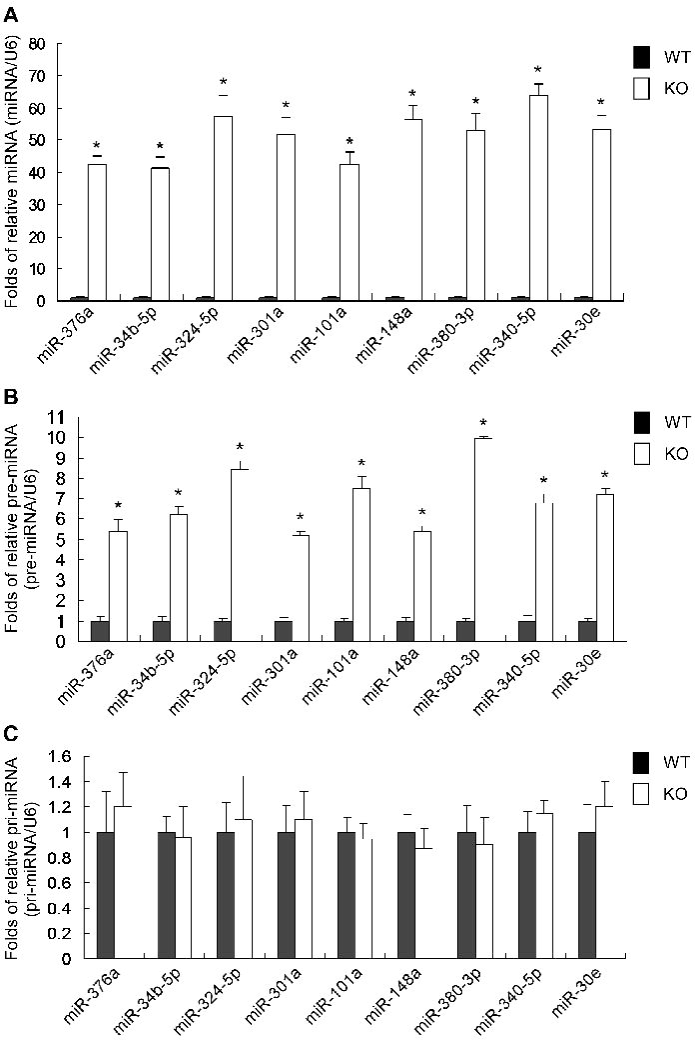
Confirmation of the 9 up-regulated miRNAs with changes over 100 folds in microarray data using qRT-PCR. (*A, B*) The changes of the miRNAs and and their pre-miRNAs in qPCR showing concordance with the microarray data. (*C*) No change of the pri-miRNAs between the WT mice and the KO mice. The relative miRNA or their precursors (miR-376a/U6) level in the WT mice was normalized as “1”. \**P* < 0.001, n=9. All experiments were performed with a biological repeat of 3 individual mice and a technical replicate for three times.

**Figure 4.**
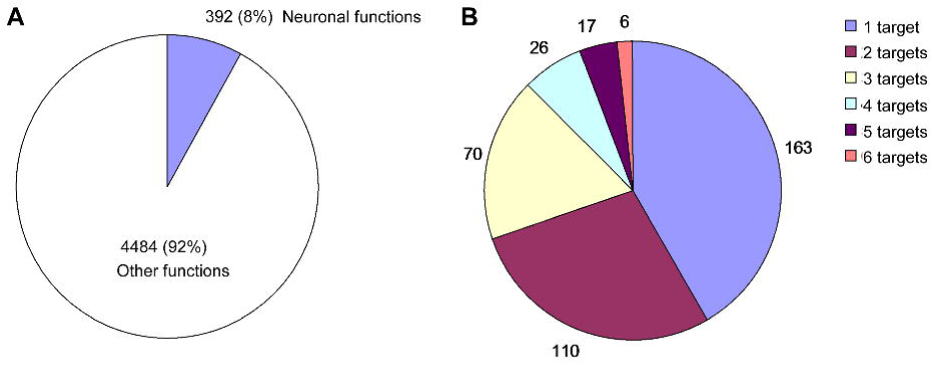
Prediction of the protein-coding genes targeted by the 9 up-regulated miRNAs with changes over 100 folds in microarray data. (*A*) Gene Ontology (GO) analysis showed that 392 (8%) target genes were associated with neuronal functions. (*B*) One to 6 targets were predicted in the 392 target genes with neuronal functions.

**Figure 5.**
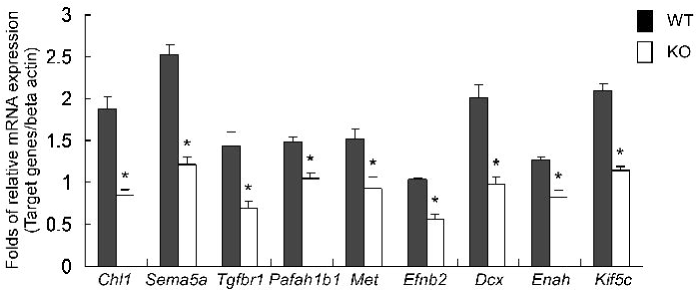
The qRT-PCR confirmation of the 9 target genes with highly expressed in the hippocampus. The relative mRNA levels (*target genes*/*beta actin*) miRNA in the KO mice were significant lower than those in the WT mice. \**P* < 0.001, n=9. All experiments were performed with a biological repeat of 3 individual mice and a technical replicate for three times.

**Figure 6.**
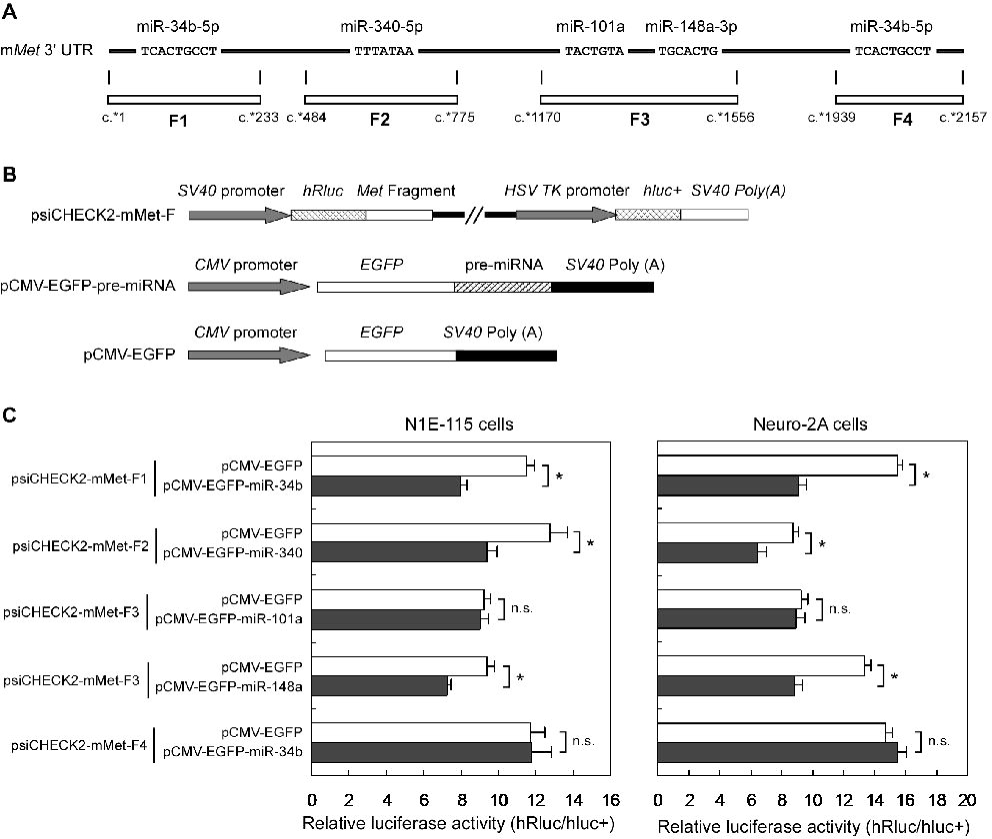
Identification of miRNA targets in the m*Met* 3′ UTR. (*A*) Predicted miRNA targets in the m*Met* 3′ UTR. The nucleotides indicate the seeds of miRNAs. F1, F2, F3 and F4 represent the fragments used for producing luciferase constructs. (*B*) Schematic of the reporter constructs used to evaluate the role of complementarity between miRNAs and the m*Met* 3′ UTR. (*C*) Luciferase activity assays of mouse Ne1-115 cells and Neuro-2A cells cotransfected with a serial of psiCHECK2-mMet-F constructs and a serial of pCMV-EGFP-pre-miRNA constructs, respectively. The *Renilla* luciferase activity was normalized to the firefly luciferase activity. \**P* < 0.01, n = 10; n.s. indicates no significant difference.

FMRP is expressed within neurons throughout the brain, with high levels in hippocampus and cerebellum (Feng, Gutekunst, Eberhart, Yi, Warren and Hersch 1997). Since FMRP is involved in synapse formation which is important for learning and memory in the hippocampus (Braun and Segal 2000; Guo et al. 2011; Jarrard 1993; Thompson 1986), we intend to investigate the regulatory role of FMRP only in the hippocampus. Our results show a development-dependant dynamic expression of *Fmr1* mRNA in mouse hippocampus with a peak at P7, suggesting an important role of FMRP in the hippocampal development and function. The P7 KO mouse, corresponding to the beginning of human full-term infant (Romijn et al. 1991), is accepted as a model for the FXS (Bilousova et al. 2009), which is the reason why we investigated the miRNA expression profile in the hippocampus of P7 *Fmr1* KO mouse in this study.

Using a genome-wide miRNA microarray analysis, we observed that the levels of 68 miRNAs significantly altered (*P* < 0.01) in the KO mice comparing to that in the wildtype mice, suggesting a critical role of FMRP in selectively regulating miRNA expression. Furthermore, using qRT-PCR experiment, we demonstrate that the highest upregulation (over 100 folds) of the 9 miRNAs in the hippocampus of *Fmr1* KO mice are resulted from miRNA processing of pri-miRNA into pre-miRNA, suggesting that FMRP should be involved in pri-miRNA processing in the nucleus. Because it is well known that FMRP shows a nuclear localization and participates in transferring mRNA from nucleus to cytoplasm (Eberhart, Malter, Feng and Warren 1996; Feng, Gutekunst, Eberhart, Yi, Warren and Hersch 1997; Fridell et al. 1996). In the cytoplasm, FMRP is also involved in miRNAs maturation and the control of target gene translation by targeting with miRNAs or by interacting with the Dicer and Argonaute proteins (Duan and Jin 2006; Jin, Zarnescu, Ceman, Nakamoto, Mowrey, Jongens, Nelson, Moses and Warren 2004). In the nucleus, a protein complex including the nuclear RNase III Drosha initiates miRNA processing (Davis et al. 2008; Han et al. 2004; Lee et al. 2003), but no description of the involvement of FMRP in the miRNA maturation in the nucleus has been reported till now. Therefore, our results suggest a new role of FMRP in regulating miRNA processing in the nucleus, which may be associated with the protein complex including Drosha. Future studies are needed to demonstrate this speculation.

Since miRNAs are involved in diverse biological processes by regulating the expressions of numerous protein-coding genes (Lewis et al. 2005; Lewis et al. 2003), we think that those altered miRNAs in the KO mice might also contribute to the abnormalities in hippocampal development and brain functions by regulating their target genes. Bioinformatic analysis showed that about 5000 genes were potentially targeted by the 9 up-regulated miRNAs and 392 of them were associated with neuronal functions. Using qRT-PCR, we found that the expressions of 9 hippocampal-expressed target genes were down-regulated in the KO mice. To determine the downregulation of the target gene resulted from the increased miRNAs, we carried out a luciferase assay to investigate the multiple targets for miR-34b-5p, miR-340-5p, miR-101a, miR-148a-5p in *Met* 3′ UTR and found that 3 of the 5 targets in the mouse *Met* 3′ UTR were functional in *vitro*, suggesting that the expression of *Met* might be co-regulated by these miRNAs in *vivo*, which is evidenced by a previous study that several miRNAs including miR-34b could negatively regulate MET expression in two lung adenocarcinomas cell lines (Migliore et al. 2008). Other results also showed that MET signaling is associated with brain functions (Judson et al. 2009; Martins et al. 2011), implicating that those up-regulated miRNAs might play a critical role in FXS by interacting with *MET* 3′ UTR. Based on the experimental identification of the miRNA targets in mouse *Met* 3′ UTR, we speculate that most potential miRNA targets in other genes should also be functional, suggesting that a complex FMRP-miRNA-mediated network could contribute to the biological role of FMRP in the hippocampus.

Together, this study suggests that FMRP is involved in the pri-miRNA processing in the nucleus and the FMRP-miRNA-mediated selective downregulation of their target genes in *Fmr1* KO mice in which the lack of FMRP might contribute to the brain abnormalities. Therefore, the altered miRNAs may be used as markers for human FXS, since most FXS cases have a very low level of FMRP that are caused by unstable expansion of the CGG trinucleotide repeat in the promoter region of the *FMR1* gene (Feng, Zhang, Lokey, Chastain, Lakkis, Eberhart and Warren 1995; Kremer, Pritchard, Lynch, Yu, Holman, Baker, Warren, Schlessinger, Sutherland and Richards 1991). Of note, our findings may provide the potential role of miRNAs in the diagnosis of FXS and give further insight in the molecular mechanism of the development of FXS, with possibility for future therapeutic targets.

## Methods

### Animals

The FVB *Fmr1* knockout mice and the FVB wildtype mice were kindly gifted by Dr. Ben A. Oostra (Erasmus University, Netherland). The animals were fed and reproduced at The Experimental Animal Center of Guangzhou Medical University with a routine animal care. Only male mice were used for the studies. This study was approved by The Ethical Committee for the use of experimental animals of Guangzhou Medical University.

### RNA isolation, RT-PCR and qRT-PCR

Total RNA was extracted from the mouse hippocampus using Trizol reagent (Invitrogen, USA) according to manufacturer’s instructions. To remove the DNA contamination, RNA samples were treated with *DNase* I (Fermentas, USA) at 37°C for 1 hr and underwent a second round of phenol-chloroform extraction and ethanol precipitation. The concentration and purity of total RNA fractions were determined by OD260/280 reading, and the qualities were assessed on 1.0% agarose gel.

The first strand of cDNA was synthesized using 2 μg of total RNA and oligo18T primer with RevertAid® premium first strand cDNA synthesis kit (Fermentas, USA) according to the manufacturer’s protocol. The primers for RT-PCR and qRT-PCR were listed in **Table S6**. RT-PCR analyses were performed using DreamTaq PCR Master Mix (Thermo-Scientific, USA) following the cycling reactions: step 1, 94°C for 5 min; step 2, 94°C for 30 sec; step 3, n°C for 30 sec (the exact temperature condition for each reaction was listed in **Table S6**); step 4, 72°C for 40 sec; step 5, repeat steps 2-4 for 28 times; and step 6, 72°C for 7 min. PCR products were assessed on 1.0% agarose gels. The qPCR (real-time PCR) was performed using Thunderbird SYBR qPCR Mix (Toyobo, Japan) and Rotor-Gene^TM^ Q instrument (Qiagen, Germany) according to the manufacturer’s protocols. All of the cDNA samples were diluted to 200 ng for qPCR analysis. The level of house-keeping gene *β-actin* was used as an endogenous control for normalization. The relative quantification of mRNA levels was determined using the ^ΔΔ^CT method (Schmittgen and Livak 2008).

### MicroRNA microarray and data analysis

MicroRNA microarrays were carried out using miRNA Microarray System with miRNA Complete Labeling and Hyb Kit (5190-0456) by Agilent Technologies (http://www.agilent.com). Oligonucleotide arrays were printed with trimer oligonucleotide probes (antisense to miRNAs) specific for 1080 mouse miRNAs using Agilent Microarray Hybridization Chamber Kit (G2534A), and miRNA expression profiling was performed and analyzed as previously described (Chan et al. 2005; Nunez-Iglesias et al. 2010). Briefly, total RNAs were isolated from the hippocampus of 3 *Fmr1* KO mice and 3 WT mice using Trizol reagent (Invitrogen, USA) according to the manufacturer’s instructions. The samples were purified by two additional chloroform extractions after the final extraction step in the Trizol protocol to ensure the removal of residual Trizol reagent. The RNAs were labeled with Cyanine 3-pCp and then hybridized to the miRNA array. To ensure accuracy of the hybridizations, each labeled RNA sample was hybridized with three separate membranes.

Hybrid signals were extracted and analyzed by Agilent Feature Extraction software (v10.7). The data were normalized and removed the unreliable signals by Agilent GeneSpring software. ANOVA’s *t* test was used to analyze the differentially expressed miRNAs among each group.

### The qRT-PCR procedures of miRNAs, pre-miRNAs and pri-miRNAs

The qRT-PCR experiments for miRNAs, pre-miRNAs and pri-miRNAs were performed according to previous descriptions (Liu et al. 2012; Schmittgen et al. 2004). Briefly, the frist-strand cDNAs of miRNAs were synthesized from 500 ng total RNA using TaqMan MicroRNA Reverse Transcription Kit (Applied Biosystems, Germany). Bulge-loop™ miRNA qRT-PCR Primer Sets designed by Guangzhou RiboBio (RiboBio, China) contained one RT primer and qPCR primer pairs for 9 miRNAs and U6 rRNA (as a control). For the reverse transcription of pre-miRNAs and pri-miRNAs, a 1 µg aliquot of total RNA was reverse transcribed to cDNA using gene-specific primer mix (listed in Table S6) with ThermoScrip RT-PCR System (invitrogen, USA) according to the manufacturer’s protocol. The qPCR was carried out with SYBR Green Real-Time PCR Master Mix Kit (Toyobo, Japan) in a 20 µl reaction volume (10 µl Sybr green mix, 200 mM forward and reverse primers, 200 ng cDNA) on a Rotor-Gene Q instrument (Qiagen, Germany) according to the indicated manufacturer’s protocol. The PCR procedure was as follow: step 1, 95°C for 60 s; step 2, 45 cycles of 95°C for 10 s, 60°C for 20 s, 70°C for 10 s. Relative miRNAs expression was calculated using a standard curve. All data were analyzed using Applied Biosystems Prism software and the ΔΔCt method.

### Prediction of miRNA targets and biological annotation analysis

To improve the accuracy of the miRNA target prediction, three computer-aided algorithms including TargetScan (Lewis, Shih, Jones-Rhoades, Bartel and Burge 2003), miRanda (John et al. 2004) and PicTar (Krek et al. 2005) were used in this study. The online website of these computer programs were as follows: http://www.targetscan.org/ for TargetScan, http://www.microrna.org/ for miRanda and http://pictar.bio.nyu.edu/ for PicTar. For online prediction, the 3′ UTRs of all annotated genes were restricted for the prediction of miRNA targets. Mouse 3′-UTR sequences were downloaded form the UCSC genome browser (Kuhn et al. 2007). Biological annotation analysis for the targeted genes was performed using the MAS 3.0 Program (http://bioinfo.capitalbio.com/mas3/), followed by Gene Ontology (GO) analysis (Tao et al. 2007).

### DNA constructs

*Met* 3′ UTR fragments (F1 to F4) were amplified from the mouse genomic DNA using primers listed in Table S4. The PCR products containing F1 to F4 were respectively cloned into the psiCHECK-2 (Promega) at *Xho* I and *Sal* I sites to produce a serial of recombinant constructs psiCHECK2-mMet-F (Fig. 6B). To clone the genomic DNA sequences containing pre-miRNA coding sequences (CDSs), the sequences ranging from about 100 bp upstream and downstream of the pre-miRNA CDSs were amplified from the genomic DNA by using primers shown in Table S6. The PCR fragments were cloned into the pCMV-EGFP to generate recombinant constructs pCMV-EGFP-miRNAs (Fig. 6B). The pre-miRNA CDSs were inserted directly downstream of the translation stop codon of the GFP CDS and upstream of the SV40 Poly (A) region to ensure pre-miRNA CDSs are only transcribed with the GFP, instead of being translated. All constructs were confirmed by sequencing.

### Transfection and luciferase assay

Mouse neuroblastoma N1E-115 and Neuro-2A cells were cultured in DMEM (GIBCO, USA) supplemented with 10% FBS (GIBCO, USA) at 37°C in a 5% CO_2_ atmosphere in 96-well plates. Cells were transfected with 200 ng of the luciferase reporter vector and 200 ng of the miRNA-expressing vector or the control vector (pCMV-EGFP) in a final volume of 0.2 ml using Lipofectamine 2000 (Invitrogen, USA). Firefly and Renilla luciferase activities were measured consecutively using Dual-Luciferase Reporter Assay System (Promega, USA) 48 hr after transfection. For each plasmid, at least 10 transfections (three experiments involving three independent plasmid preparations) were carried out. The activities of *Renilla* luciferase and firefly luciferase were detected by a GloMax 20/20 Luminometer (Promega, USA) according to the manufacturer’s instructions. The relative activities (hRluc/hluc+) are expressed as mean±SEM.

### Western blotting analysis

Protein extracts were prepared from mouse hippocampus using T-PER Tissue Protein Extraction Reagent (Pierce, USA). The protein concentrations in the fractions were determined by the bicinchoninic acid (BCA) assay. Purified extracts were subjected to SDS polyacrylamide gel electrophoresis and then transferred to polyvinylidene difluoride (PVDF) membranes (Millipore, Bedford, MA). The membranes were incubated at room temperature for 4 hours in 10% milk blott solution (50 mM Tris, 80 mM NaCl, 10% skimmed milk power, 0.1% Tween 20, pH 8.0). After being rinsed with TBST, the membranes were incubated with a polyclonal anti-FMRP antibody (Abcam, Cambridge, MA) and a polyclonal anti-GAPDH antibody (Abcam, Cambridge, MA) for 16 hours at 4°C. A 1:4000 dilution of Peroxidase Streptavidin (Jackson ImmunoResearch, PA) was made in TBST and the membranes were incubated at room temperature for 1 hour with shaking. After washing 3 times in TBST, Primary antibody binding sites were visualized using enhanced chemiluminescence.

### Statistical analysis

All experimental data were analyzed using ANOVA and Tukey′s *t* test. The significance level for all statistical tests was 0.05.

## Acknowledgements

We are indebted to Dr. Ben A. Oostra (Erasmus University, Netherland) for kindly providing the FVB *Fmr1* knockout mice and the FVB wildtype mice. We are grateful to the He Shanheng Charity Foundation for contributing to the development of this institute. YSL was supported by National Natural Science Foundation of China (81371436 and 31070928), Guangzhou Scholar Project (10A011G), and Scientific Research of Guangzhou Municipal Colleges and Universities (10A211). YHY was supported by National Natural Science Foundation of China (81171073 and 30870876). WPL was supported by National Natural Science Foundation of China (81271434).

---

**Figure S1.** The mRNAs of partial miRNA target genes present in the mouse hippocampus. These data of the mRNA levels determined by in situ hybridization were downloaded from the public data from Allen Brain Atlas Database (http://www.brain-map.org/).

## References

Antar LN, Dictenberg JB, Plociniak M, Afroz R, and Bassell GJ. 2005. Localization of FMRP-associated mRNA granules and requirement of microtubules for activity-dependent trafficking in hippocampal neurons. Genes Brain Behav 4: 350–359.

Ashley CT, Jr., Wilkinson KD, Reines D, and Warren ST. 1993. FMR1 protein: conserved RNP family domains and selective RNA binding. Science 262: 563–566.

Bailey DB, Jr., Hatton DD, Skinner M, and Mesibov G. 2001. Autistic behavior, FMR1 protein, and developmental trajectories in young males with fragile X syndrome. J Autism Dev Disord 31: 165–174.

Bilousova TV, Dansie L, Ngo M, Aye J, Charles JR, Ethell DW, and Ethell IM. 2009. Minocycline promotes dendritic spine maturation and improves behavioural performance in the fragile X mouse model. J Med Genet 46: 94–102.

Blackwell E, Zhang X, and Ceman S. 2010. Arginines of the RGG box regulate FMRP association with polyribosomes and mRNA. Hum Mol Genet 19: 1314–1323.

Blonden L, van ‘t Padje S, Severijnen LA, Destree O, Oostra BA, and Willemsen R. 2005. Two members of the Fxr gene family, Fmr1 and Fxr1, are differentially expressed in Xenopus tropicalis. Int J Dev Biol 49: 437–441.

Braun K and Segal M. 2000. FMRP involvement in formation of synapses among cultured hippocampal neurons. Cereb Cortex 10: 1045–1052.

Chan JA, Krichevsky AM, and Kosik KS. 2005. MicroRNA-21 is an antiapoptotic factor in human glioblastoma cells. Cancer Res 65: 6029–6033.

Chelly J and Mandel JL. 2001. Monogenic causes of X-linked mental retardation. Nat Rev Genet 2: 669–680.

Darnell JC and Klann E. 2013. The translation of translational control by FMRP: therapeutic targets for FXS. Nat Neurosci 16: 1530–1536.

Darnell JC, Van Driesche SJ, Zhang C, Hung KY, Mele A, Fraser CE, Stone EF, Chen C, Fak JJ, Chi SW et al. 2011. FMRP stalls ribosomal translocation on mRNAs linked to synaptic function and autism. Cell 146: 247–261.

Davis BN, Hilyard AC, Lagna G, and Hata A. 2008. SMAD proteins control DROSHA-mediated microRNA maturation. Nature 454: 56–61.

De Boulle K, Verkerk AJ, Reyniers E, Vits L, Hendrickx J, Van Roy B, Van den Bos F, de Graaff E, Oostra BA, and Willems PJ. 1993. A point mutation in the FMR-1 gene associated with fragile X mental retardation. Nat Genet 3: 31–35.

Duan R and Jin P. 2006. Identification of messenger RNAs and microRNAs associated with fragile X mental retardation protein. Methods Mol Biol 342: 267–276.

Eberhart DE, Malter HE, Feng Y, and Warren ST. 1996. The fragile X mental retardation protein is a ribonucleoprotein containing both nuclear localization and nuclear export signals. Hum Mol Genet 5: 1083–1091.

Edbauer D, Neilson JR, Foster KA, Wang CF, Seeburg DP, Batterton MN, Tada T, Dolan BM, Sharp PA, and Sheng M. 2010. Regulation of synaptic structure and function by FMRP-associated microRNAs miR-125b and miR-132. Neuron 65: 373–384.

Feng Y, Gutekunst CA, Eberhart DE, Yi H, Warren ST, and Hersch SM. 1997. Fragile X mental retardation protein: nucleocytoplasmic shuttling and association with somatodendritic ribosomes. J Neurosci 17: 1539–1547.

Feng Y, Zhang F, Lokey LK, Chastain JL, Lakkis L, Eberhart D, and Warren ST. 1995. Translational suppression by trinucleotide repeat expansion at FMR1. Science 268: 731–734.

Fridell RA, Benson RE, Hua J, Bogerd HP, and Cullen BR. 1996. A nuclear role for the Fragile X mental retardation protein. EMBO J 15: 5408–5414.

Guo W, Allan AM, Zong R, Zhang L, Johnson EB, Schaller EG, Murthy AC, Goggin SL, Eisch AJ, Oostra BA et al. 2011. Ablation of Fmrp in adult neural stem cells disrupts hippocampus-dependent learning. Nat Med 17: 559–565.

Han J, Lee Y, Yeom KH, Kim YK, Jin H, and Kim VN. 2004. The Drosha-DGCR8 complex in primary microRNA processing. Genes Dev 18: 3016–3027.

Hessl D, Tassone F, Loesch DZ, Berry-Kravis E, Leehey MA, Gane LW, Barbato I, Rice C, Gould E, Hall DA et al. 2005. Abnormal elevation of FMR1 mRNA is associated with psychological symptoms in individuals with the fragile X premutation. Am J Med Genet B Neuropsychiatr Genet **139B**: 115–121.

Hirst M, Grewal P, Flannery A, Slatter R, Maher E, Barton D, Fryns JP, and Davies K. 1995. Two new cases of FMR1 deletion associated with mental impairment. Am J Hum Genet 56: 67–74.

Huber KM, Gallagher SM, Warren ST, and Bear MF. 2002. Altered synaptic plasticity in a mouse model of fragile X mental retardation. Proc Natl Acad Sci U S A 99: 7746–7750.

Jacquemont S, Hagerman RJ, Hagerman PJ, and Leehey MA. 2007. Fragile-X syndrome and fragile X-associated tremor/ataxia syndrome: two faces of FMR1. Lancet Neurol 6: 45–55.

Jarrard LE. 1993. On the role of the hippocampus in learning and memory in the rat. Behav Neural Biol 60: 9–26.

Jin P, Zarnescu DC, Ceman S, Nakamoto M, Mowrey J, Jongens TA, Nelson DL, Moses K, and Warren ST. 2004. Biochemical and genetic interaction between the fragile X mental retardation protein and the microRNA pathway. Nat Neurosci 7: 113–117.

John B, Enright AJ, Aravin A, Tuschl T, Sander C, and Marks DS. 2004. Human MicroRNA targets. PLoS Biol 2: e363.

Judson MC, Bergman MY, Campbell DB, Eagleson KL, and Levitt P. 2009. Dynamic gene and protein expression patterns of the autism-associated met receptor tyrosine kinase in the developing mouse forebrain. J Comp Neurol 513: 511–531.

Krek A, Grun D, Poy MN, Wolf R, Rosenberg L, Epstein EJ, MacMenamin P, da Piedade I, Gunsalus KC, Stoffel M et al. 2005. Combinatorial microRNA target predictions. Nat Genet 37: 495–500.

Kremer EJ, Pritchard M, Lynch M, Yu S, Holman K, Baker E, Warren ST, Schlessinger D, Sutherland GR, and Richards RI. 1991. Mapping of DNA instability at the fragile X to a trinucleotide repeat sequence p(CCG)n. Science 252: 1711–1714.

Kuhn RM, Karolchik D, Zweig AS, Trumbower H, Thomas DJ, Thakkapallayil A, Sugnet CW, Stanke M, Smith KE, Siepel A et al. 2007. The UCSC genome browser database: update 2007. Nucleic Acids Res 35: D668–673.

Lee Y, Ahn C, Han J, Choi H, Kim J, Yim J, Lee J, Provost P, Radmark O, Kim S et al. 2003. The nuclear RNase III Drosha initiates microRNA processing. Nature 425: 415–419.

Lewis BP, Burge CB, and Bartel DP. 2005. Conserved seed pairing, often flanked by adenosines, indicates that thousands of human genes are microRNA targets. Cell 120: 15–20.

Lewis BP, Shih IH, Jones-Rhoades MW, Bartel DP, and Burge CB. 2003. Prediction of mammalian microRNA targets. Cell 115: 787–798.

Liu N, Chen NY, Cui RX, Li WF, Li Y, Wei RR, Zhang MY, Sun Y, Huang BJ, Chen M et al. 2012. Prognostic value of a microRNA signature in nasopharyngeal carcinoma: a microRNA expression analysis. Lancet Oncol 13: 633–641.

Lu R, Wang H, Liang Z, Ku L, O’Donnell W T, Li W, Warren ST, and Feng Y. 2004. The fragile X protein controls microtubule-associated protein 1B translation and microtubule stability in brain neuron development. Proc Natl Acad Sci U S A 101: 15201–15206.

Martins GJ, Shahrokh M, and Powell EM. 2011. Genetic disruption of Met signaling impairs GABAergic striatal development and cognition. Neuroscience 176: 199–209.

Mercaldo V, Descalzi G, and Zhuo M. 2009. Fragile X mental retardation protein in learning-related synaptic plasticity. Mol Cells 28: 501–507.

Michel CI, Kraft R, and Restifo LL. 2004. Defective neuronal development in the mushroom bodies of Drosophila fragile X mental retardation 1 mutants. J Neurosci 24: 5798–5809.

Migliore C, Petrelli A, Ghiso E, Corso S, Capparuccia L, Eramo A, Comoglio PM, and Giordano S. 2008. MicroRNAs impair MET-mediated invasive growth. Cancer Res 68: 10128–10136.

Nunez-Iglesias J, Liu CC, Morgan TE, Finch CE, and Zhou XJ. 2010. Joint genome-wide profiling of miRNA and mRNA expression in Alzheimer’s disease cortex reveals altered miRNA regulation. PLoS One 5: e8898. doi: 10.1371/journal.pone.0008898.

O’Donnell WT and Warren ST. 2002. A decade of molecular studies of fragile X syndrome. Annu Rev Neurosci 25: 315–338.

Romijn HJ, Hofman MA, and Gramsbergen A. 1991. At what age is the developing cerebral cortex of the rat comparable to that of the full-term newborn human baby? Early Hum Dev 26: 61–67.

Schmittgen TD, Jiang J, Liu Q, and Yang L. 2004. A high-throughput method to monitor the expression of microRNA precursors. Nucleic Acids Res 32: e43.

Schmittgen TD and Livak KJ. 2008. Analyzing real-time PCR data by the comparative C(T) method. Nat Protoc 3: 1101–1108.

Sidorov MS, Auerbach BD, and Bear MF. 2013. Fragile X mental retardation protein and synaptic plasticity. Mol Brain 6: 15. doi: 10.1186/1756-6606-6-15.

Siomi H, Choi M, Siomi MC, Nussbaum RL, and Dreyfuss G. 1994. Essential role for KH domains in RNA binding: impaired RNA binding by a mutation in the KH domain of FMR1 that causes fragile X syndrome. Cell 77: 33–39.

Tao Y, Sam L, Li J, Friedman C, and Lussier YA. 2007. Information theory applied to the sparse gene ontology annotation network to predict novel gene function. Bioinformatics 23: i529–538.

Thompson RF. 1986. The neurobiology of learning and memory. Science 233: 941–947.

Trottier Y, Imbert G, Poustka A, Fryns JP, and Mandel JL. 1994. Male with typical fragile X phenotype is deleted for part of the FMR1 gene and for about 100 kb of upstream region. Am J Med Genet 51: 454–457.

Verheij C, Bakker CE, de Graaff E, Keulemans J, Willemsen R, Verkerk AJ, Galjaard H, Reuser AJ, Hoogeveen AT, and Oostra BA. 1993. Characterization and localization of the FMR-1 gene product associated with fragile X syndrome. Nature 363: 722–724.

Verkerk AJ, Pieretti M, Sutcliffe JS, Fu YH, Kuhl DP, Pizzuti A, Reiner O, Richards S, Victoria MF, Zhang FP et al. 1991. Identification of a gene (FMR-1) containing a CGG repeat coincident with a breakpoint cluster region exhibiting length variation in fragile X syndrome. Cell 65: 905–914.

Zang JB, Nosyreva ED, Spencer CM, Volk LJ, Musunuru K, Zhong R, Stone EF, Yuva-Paylor LA, Huber KM, Paylor R et al. 2009. A mouse model of the human Fragile X syndrome I304N mutation. PLoS Genet 5: e1000758. doi: 10.1371/journal.pgen.1000758.

